# From eye anatomy to navigation: a biologically accurate model of bees’ polarisation vision

**DOI:** 10.64898/2026.06.01.729196

**Authors:** George E Kolyfetis, Evripidis Gkanias, Athil A Aliyam Veetil Zynudheen, Vun Wen Jie, Giovanni C Galizia, Emily Baird, Barbara Webb, James J Foster

## Abstract

Skylight polarisation patterns provide a critical navigational cue for many insects. Bees perceive these patterns through specialised ommatidia in the dorsal rim area of their compound eyes, enabling them to estimate the sun’s direction and navigate between food sources and the hive. Although polarisation-based navigation has been extensively studied behaviourally, computational models that link DRA anatomy with navigational performance are lacking. Here, we simulate polarisation vision in honeybees (*Apis mellifera*) and bumblebees (*Bombus terrestris*) using real sky polarisation images to capture biologically relevant skylight properties. Our biologically grounded simulation incorporates species-specific DRA anatomy, including ommatidial optical axis directions, photoreceptor receptive fields, and microvillar orientations. We evaluate navigational accuracy and consistency across sun elevations under two distinct, potentially complementary navigational models: the matched filter, which requires scanning across body orientations to identify the solar axis, and the vector-sum model, which generates instantaneous sun azimuth estimates from a single body orientation, making it independent of active scanning. Matched filter errors in estimating solar axis are below 5° across most sun elevations and in both species. Absolute errors in the vector-sum model are lower for honeybees than bumblebees (median ∼10° and ∼30°, respectively), reflecting differences in DRA anatomy, particularly viewing direction and microvillar arrangement. Both models allow stable course control across most sun elevations in both species, yet the matched filter, being limited to solar axis alignment, only enables positive or negative phototaxis. Overall, this work provides a mechanistic and comparative framework based on realistic DRA anatomy to study polarisation-based navigation, generating testable predictions for insect navigation under natural sky conditions.

**Author Summary:** Many insects, including bees, navigate with the help of skylight polarisation patterns which hold information about the sun’s position even when it is not visible. Bees detect these patterns through the dorsal rim area (DRA) of their complex eyes. How differences in DRA anatomy between bee species translate into differences in navigational ability has remained unclear. Here, we built a biologically realistic simulation of polarisation vision in honeybees and bumblebees. We used real sky images to examine what polarisation information is available to each species. We then tested two models of sun position estimation based on the polarisation pattern: one that requires the bee to actively scan the sky, and one that generates an instantaneous estimate from a single body orientation. In both species, both models show that accurate sun position estimation and stable navigation are possible using just polarisation information under a wide range of sun elevations. Differences in navigational performance between honeybees and bumblebees arise because the two DRAs look at different parts of the sky. Our results provide a robust framework for understanding how DRA anatomy shapes polarisation-based navigation in bees.

## Introduction

The ability to direct behaviour and navigate in a changing visual environment is crucial for the survival and fitness of many insect species. The sun serves as one reliable navigational cue and many insect species track its position to orient themselves^1–5^. However, when direct information about the sun’s azimuth is ambiguous, e.g., when it is covered by clouds, the skylight polarisation pattern that the sun creates in the sky provides insects with a robust source of directional information, aiding them in finding their way home, a food source or maintaining a steady course^6–15^. The process by which polarisation information gets transduced into a navigational cue to create this ‘polarisation compass’ represents a complex biological phenomenon that has not yet been modelled with both realistic sky properties and the anatomical detail of insect eyes.

The primary input for this polarisation compass, which helps insects track the position of the sun, is partially polarised light coming from the sky. Skylight polarisation patterns are the result of linearly polarised light, generated in the sky dome by scattering processes as sunlight passes the Earth’s atmosphere, and the topography of these patterns is determined by the sun’s location. The primary angle of oscillation of the light waves at any given point in the sky determines the angle of linear polarisation (AoLP) and the proportion of light oscillating at the AoLP defines the degree of linear polarisation (DoLP)^16,17^. Although polarisation patterns in the sky can be predicted from physical principles, in practice various causes contribute to produce high spatial complexity^18^, not fully captured in previous realistic but simplified simulations^19–21^.

Most insects, including bees, detect polarised light through an upward-viewing region of their compound eyes, called the dorsal rim area (DRA)^22–24^. The DRA ommatidia exhibit unique anatomical and functional features, crucial for the detection of polarised light. As a result of their untwisted rhabdoms, their photoreceptors have high polarisation sensitivity (PS) and their receptive fields (RFs) are extended compared to ommatidia from other parts of the eye. This is possibly an optimization to sample large sky regions and smooth out disturbances (e.g., due to clouds) of the polarisation pattern^22,25,26^. In DRA photoreceptors, the microvilli, i.e., the membranes containing photosensitive opsin molecules, are arranged orthogonally, making the two sets of photoreceptors of each ommatidium sensitive to two perpendicular AoLP^27^. Responses from the two photoreceptor sets are compared by integrative POL-neurons that show a sinusoidal activation pattern across AoLPs^28–30^. The AoLP of maximum activation, corresponds to the orientation angle of one of the two photoreceptor sets. With reference to the body axis, the microvilli are not uniformly oriented across the DRA ommatidia of many insect species. Rather, their orientation gradually rotates from the anterior to the posterior part of the DRA creating a fan-shaped arrangement of microvilli across the DRA^31–37^. These anatomical features shape how polarisation information is sampled and integrated, constraining how accurately the sun’s azimuth can be estimated and used for navigation.

The level of accuracy of polarisation-based orientation behaviour under different sky conditions can reveal how insects process polarisation information. Behavioural experiments on the honeybee waggle dance, a behaviour that conveys the location of a food source with reference to the sun^38^, suggest errors as low as 10° under clear skies. Navigation performance is reduced, however, when animals are presented with polarisation stimuli of low DoLP^39,40^ or low intensity^41^. A restricted view of the sky containing a single AoLP is also known to elicit ambiguous, bimodal waggle dances^42^. Different hypotheses have been proposed to explain the mechanism behind bees’ systematic navigational errors. The ‘matched filter’ hypothesis suggests that the fan-shaped arrangement of DRA microvilli, i.e., the ‘filter’, plays a crucial role in estimating the solar axis accurately. Its proposed function is to act as a preset filter of polarisation directions that matches the AoLP pattern in the sky when the animal is aligned with the solar axis^32,43,44^. More specifically, bees are assumed, under this hypothesis, to scan the sky by rotating about their vertical axis to locate the position of maximum alignment with the AoLP pattern. The degree of this alignment is proportional to how well they can estimate the position of the sun, which can lead to systematic errors^32,43,44^. An alternative hypothesis suggests that such scanning of the sky is not necessary for bees to accurately estimate the sun’s azimuth. Bees may recover it instantaneously from a single body orientation while looking at the sky’s polarisation pattern (Gkanias et al., in preparation)^41,42^. These hypotheses offer a great opportunity for computational assessment and quantification as they make explicit predictions about the relationship between skylight polarisation patterns and navigational performance.

The polarisation compass was behaviourally demonstrated in honeybees and desert ants several decades ago^42,45,46^. The bumblebee polarisation compass has also gained increased attention recently, with recent studies focusing on DRA visual properties^26,47^ and polarisation-based navigational performance^48^. However, the few attempts to model the mechanism of the compass computationally have mainly centred around optimising visual parameters for insect-inspired robotics^29,30,49–52^. As a result, the functional effects of DRA anatomy on navigational accuracy remain poorly understood. In this study, we bridge the gap between eye anatomy and polarisation-based navigation to create a biologically accurate model of the DRA and polarisation compass in honeybees (*Apis mellifera*) and bumblebees (*Bombus terrestris*) (Fig.1a). By minimising assumptions about skylight properties with real polarisation sky images (Fig.1b), we quantitatively assess two distinct, though not mutually exclusive, models of sun position estimation: the scanning matched filter model^43^ and the instantaneous vector-sum model (Gkanias et al., in preparation) (Fig. 1c,d). We compare the models in the two species both in terms of accuracy and consistency of sun azimuth estimation across sun elevations. This comparative approach allows us to link species-specific DRA anatomy to navigational accuracy and to generate testable behavioural predictions under natural sky conditions.

**Fig. 1.**
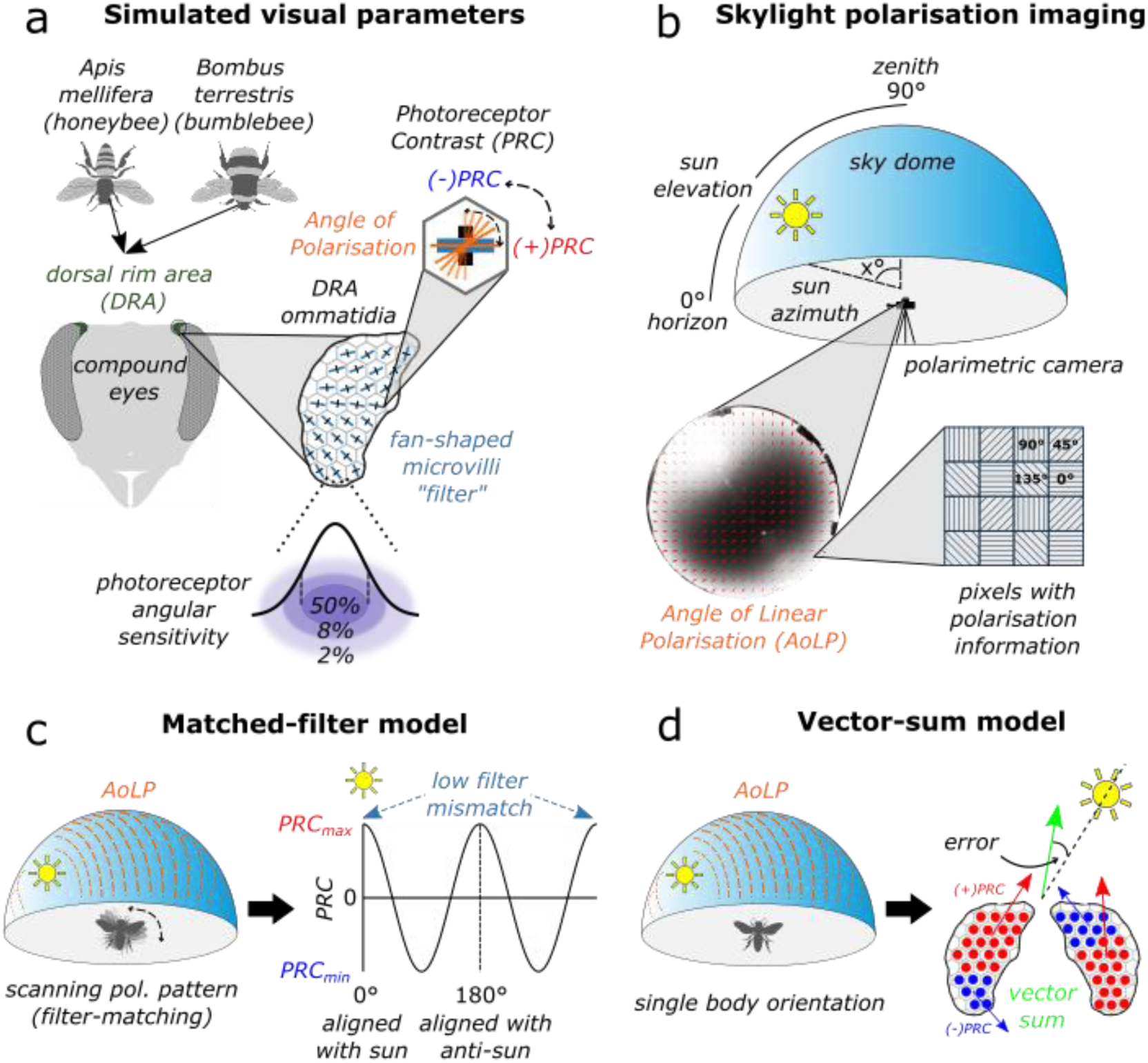
Graphical overview. a) Simulated visual parameters of the honeybee (*Apis mellifera*) and bumblebee (*Bombus terrestris*) dorsal rim area (DRA) in this study. These include the angular sensitivity of the DRA photoreceptors, the microvilli orientations of the DRA ommatidia and the photoreceptor contrasts (PRC). b) Skylight polarisation images contain per-pixel information about angle and degree of polarisation. c) The matched filter model requires sequential scanning of the polarisation pattern that produces different photoreceptor contrasts depending on body orientation. The lowest mismatch between the fan-shaped arrangement of DRA microvilli (‘filter’) and the polarisation pattern is hypothesised to be achieved when the animal is aligned with the sun and the anti-sun where the total PRC is also maximum. d) The vector-sum model can produce a sun position estimate from a single body orientation. For each DRA ommatidium a vector is constructed based on the sign and magnitude of the PRC. All individual vectors are then summed to produce a single sun position estimate.

## Methods

### Polarisation Image Processing

#### Sky imaging

An overview of our polarisation image processing is shown in Figure 2. Sky polarisation images were captured using a division-of-focal-plane polarised monochrome camera (PHX050S1-PC; LUCID Vision Labs Inc., Richmond B.C., Canada) equipped with a fisheye lens (62274; Edmund Optics Ltd., Nether Poppleton, United Kingdom). Since the daily sun trajectory across the sky influences the polarization pattern, we sampled natural sky polarization patterns across full days at different latitudes: 47°39’ (Konstanz), 43°17’ (Marseille, data from literature^53^), 37°98’ (Athens), 25°27’ (Mbombela). Specifically, for images taken in Konstanz (Germany) and Mbombela (South Africa), three images at different exposure times (low, medium, and high; Figure 2a) were acquired ten times consecutively with a short inter-acquisition delay (approximately equal to the sum of the three exposure times); the ten images per exposure level were averaged to increase signal-to-noise ratio. For images taken in Athens (Greece), a single exposure bracket was acquired and no averaging was performed.

**Figure 2.**
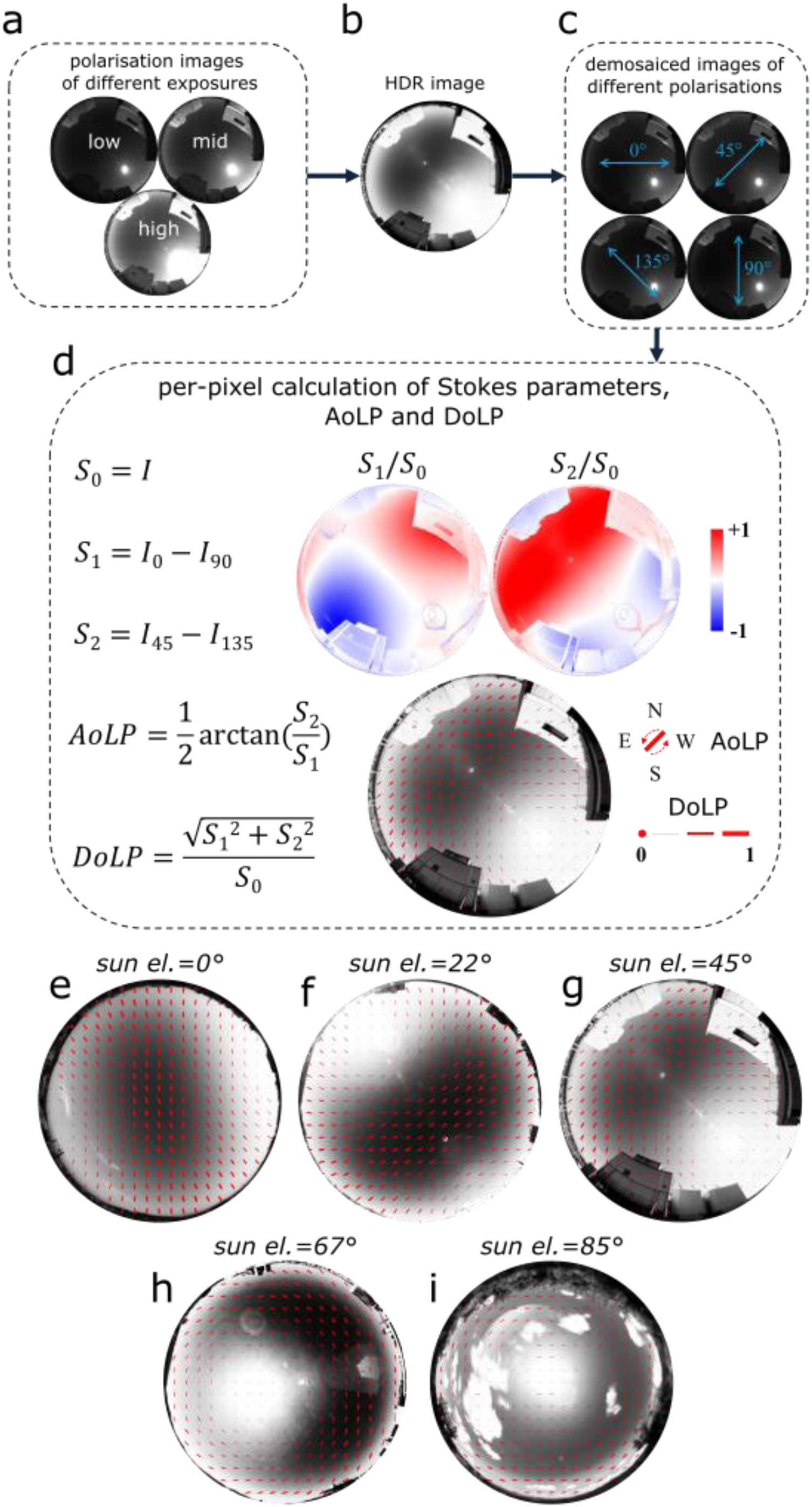
Image processing of polarisation sky images. a) Polarisation images for three exposure times (low, middle, high) were taken for each sky condition. b) The three images of different exposure times were combined into a single High Dynamic Range (HDR) image which was sigmoid scaled for visualization purposes. c) The HDR image was then decomposed (demosaiced) into four images of different polarisations, namely 0°, 45°, 90° and 135°. d) For each pixel of the HDR image, Stokes parameters, Angle of Linear Polarisation (AoLP) and Degree of Linear Polarisation (DoLP) are calculated. A positive Stokes parameters ratio (red) indicates AoLP closer to 0° (S_1_/S_0_) or 45° (S_2_/S_0_) and negative (blue) indicates AoLP closer to 90° (S_1_/S_0_) or 135° (S_2_/S_0_). e-i) Example HDR images (sigmoid-scaled) of the different sun elevations recorded, namely (e) 0°, (f) 22.5°, (g) 45°, (h) 67.5° and (i) 85° respectively. Red lines in sky images indicate the AoLP (orientation) and DoLP (thickness).

To correct for non-linearities in the camera’s response, we calibrated pixel-byte values to light intensity using reflectance standards (See Supplementary Information). Images of different exposure times were combined into a single High Dynamic Range (HDR) image (Figure 2b). The HDR image was then demosaiced into four images containing pixel intensity values that correspond to four polarisation states, 0°, 45°, 90° and 135° (https://www.sony-semicon.com/en/technology/industry/polarsens.html) (Figure 2c). From these four images, the Stokes parameters, AoLP, and DoLP were calculated for each pixel (Figure 2d).

We additionally processed images from the polarisation imaging dataset of Poughon et al. (2024), consisting of 16-bit RGB images taken in Marseille, France. These images were converted to 8-bit (0-255) float arrays, and only the blue channel was used for HDR image generation and subsequent calculation of Stokes parameters, AoLP, and DoLP, following the same calibration procedure described above. Three exposure levels were available for these images.

Camera lens spatial calibration was performed by evaluating several projection models; the azimuthal equidistant projection provided the best fit for both our imaging setup and the Poughon et al. ^53^ dataset (Supplementary Figure 2). Under this projection, radial distance in the image is linearly proportional to angular distance from the optical axis. We corrected for azimuthal equidistant distortion by normalising the image radius to 90° elevation, converting pixel distances into angular coordinates using a constant pixels-per-degree scaling.

For display purposes only, sky images were sigmoid-scaled using log-scaled linearised brightnesses, reducing visual detail across extremely bright and dark pixels while enhancing contrast across intermediate brightnesses^54^.

Polarisation images were collected across six sun elevation ranks under clear sky conditions: ≈0° (sun at horizon; 5 images), ≈23° (5 images), ≈45° (5 images), ≈66° (5 images), ≈75° (1 image), and ≈85° (sun near zenith; 2 images) (Supplementary Table 1). Example images for each condition, along with corresponding AoLP and DoLP distributions, are shown in Figure 2e–i.

The true sun azimuth was determined in two ways. When the sun was visible in the image, we used a custom-modified Python script to locate the centre of the brightest region by applying a Gaussian blur (radius = 101 pixels) to the low-exposure image, where the sun disk covered the smallest possible area^55^. For images from Konstanz and Mbombela, the first of the ten low-exposure images was used. This approach minimises errors arising from potential camera misalignment with North. When the sun was not visible, sun azimuth was obtained from www.timeanddate.com, assuming correct camera alignment with North and correcting for magnetic declination. The difference between online sun azimuth data and brightest-region predictions had a standard deviation of 1°.

#### Sky simulations

To investigate potential differences in skylight polarisation patterns and sun position estimation performance between real and simulated skies, we generated simulated sky images as azimuthal equidistant projections of the same dimensions as our polarisation camera images, following Gkanias et al. (2019) and based on established sky dome simulations^20^. Simulated images were generated for the same sun elevation conditions as the real sky images. For the analysis on simulated skies only, we added one extra image of 10° sun elevation to increase resolution at low sun elevations where the differences to real skies are more pronounced and where high navigational errors occur. For each pixel, AoLP, DoLP, and intensity (S0 Stokes parameter) were calculated, from which S1 and S2 were derived as:

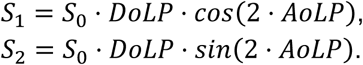

Demosaiced images for the four polarisation states (0°, 45°, 90°, 135°) were then simulated per pixel as^56^:

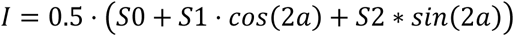

This ensured that simulated skies underwent identical processing to real-world images. The per-pixel absolute angular difference between real and simulated AoLP values was then calculated (Supplementary Figure 1).

### Dorsal Rim Area simulation

The optical axes of the individual dorsal rim area (DRA) ommatidia were obtained from micro computed-tomography (micro-CT) scans of phosphotungstic acid (PTA) stained bumblebee eyes and isolated using a localization technique^57^. The surface normals of the individual DRA ommatidia were used to obtain their optical axis. The projections of these optical axes onto the outside world were done using ray tracing^58^, with a head pitch angle similar to that found in behavioural studies during flight and landing^59^. The same pitch angle was also used for honeybees, which aligned well with the optical axes described in Labhart 1980^22^ (Supplementary Figure 2). More specifically, to reconstruct the estimated DRA FoV from Labhart, 1980 (Supplementary Figure 3), we fitted a spline through four points on a sphere (scipy.interpolate.CubicSpline), mentioned by Labhart as the limits of the DRA FoV. These points were ([azimuth,elevation]): [0°,80°]; [180°,50°]; [270°,80°] and [270°,65°].

Figure 4a*_*ii*_*,b*_*ii*_* demonstrates the receptive fields of individual DRA ommatidia clustered in two regions corresponding to the two eyes of the two species and the total FoV of the DRA. Note that the DRA of each eye has a contralateral view of the sky owing to the curvature of the DRA in the compound eye. For honeybees, angular sensitivity values of DRA photoreceptors were extracted manually from Labhart (1980) (Figure 3b therein) and a 2D spline was fitted after mirroring values across the centre, yielding the angular sensitivity function used for honeybee DRA ommatidia (Figure 4a_*i*_). Honeybee ommatidial FoVs were also modelled as 2D Gaussian kernels (σ = 2.32°), which yielded very similar results (Supplementary Figure 4). As no wide low-sensitivity area (brim) has been identified in *Bombus terrestris* ^26^, bumblebee ommatidial FoVs were modelled as 2D Gaussian kernels (σ = 2.65°; Figure 4b_*i*_), resulting in a smaller total DRA FoV (Figure 4b_*ii*_).

**Figure 3.**
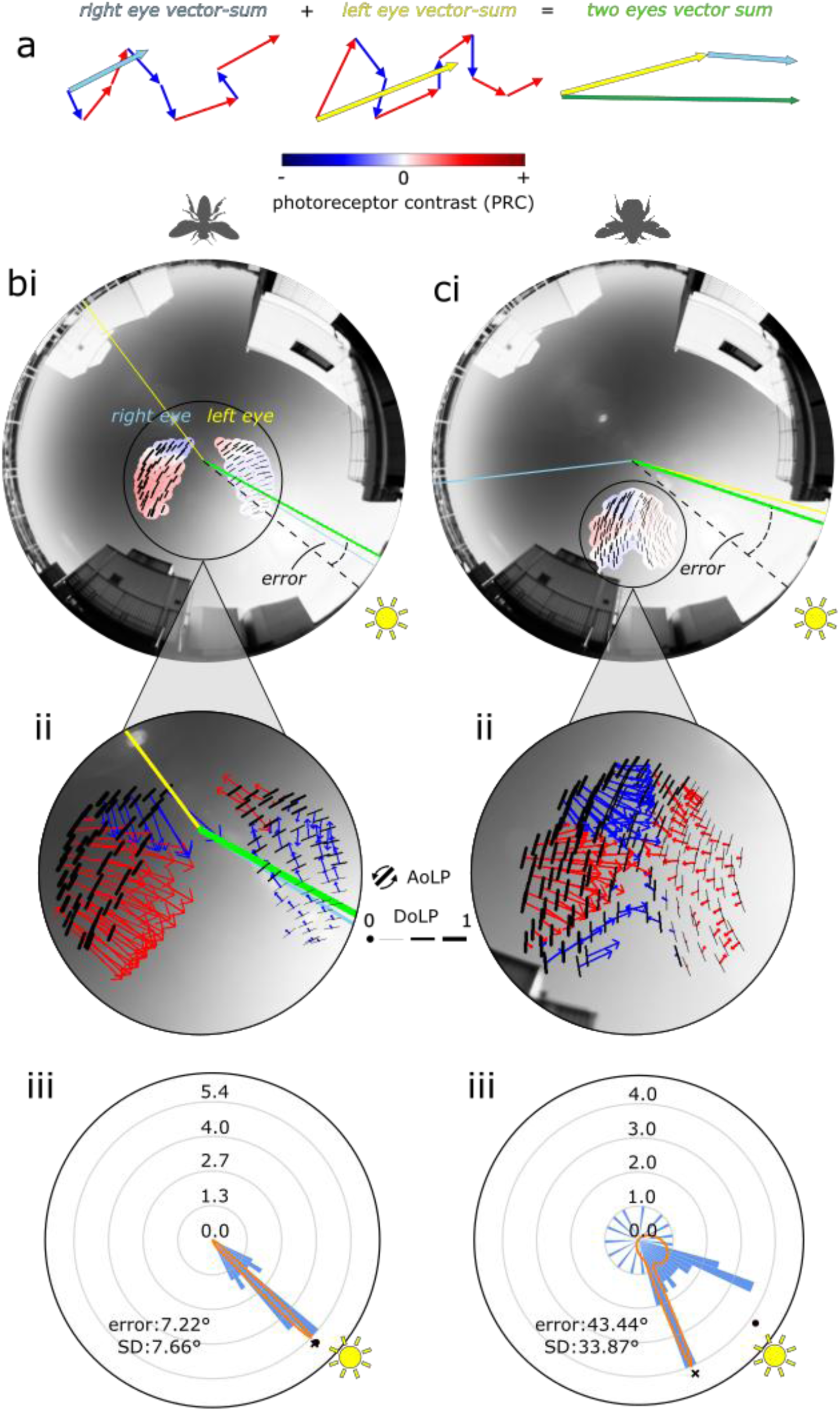
Vector-sum model for sun azimuth detection. a) Schematic of the vector-sum model. For each eye, one photoreceptor contrast (PRC) value between φ_max1_ and φ_max2_ is calculated per ommatidium and represented as a vector. Red arrows indicate positive PRC, blue arrows indicate negative PRC. The vector length (response magnitude) is shown as arrow length. The vectors are summed across ommatidia and the two eyes (light blue and yellow arrows). The resulting vector-sum (green arrow) represents the total sun azimuth estimation. b-c) (i) Vector-sum model applied to polarisation images for the honeybee (b) and bumblebee (c) DRA. Small circles indicate individual ommatidia 100%-50% angular sensitivity area (Full Width at Half Maximum) and are coloured red for positive PRC and blue for negative PRC. Black lines indicate AoLP (orientation) and DoLP (thickness) viewed by the individual ommatidia. Dashed line indicates the sun azimuth, yellow line indicates the left eye’s sun azimuth estimate, light blue line indicates right eye’s sun azimuth estimate (contralateral FoVs) and green line indicates total vector-sum estimate. The error is defined as the angular distance between sun azimuth and total vector-sum estimate. (ii) Visualization of individual ommatidia PRC vectors. The vectors are added to produce the total vector-sum sun azimuth estimate. (iii) To summarize the vector-sum model performance for a given sun position, each image is sampled 72 times across 360° (72 samples, 5° apart), and a sun azimuth estimate is calculated for each sample. A mean absolute error and a standard deviation (SD) are computed and a mixture of two von Mises distributions (orange line) is fitted to the data. The rose histogram depicts the square root of the frequency of counts (sun position estimates). The main von Mises mode is indicated by a black dot and the secondary von Mises mode by a black cross.

**Figure 4.**
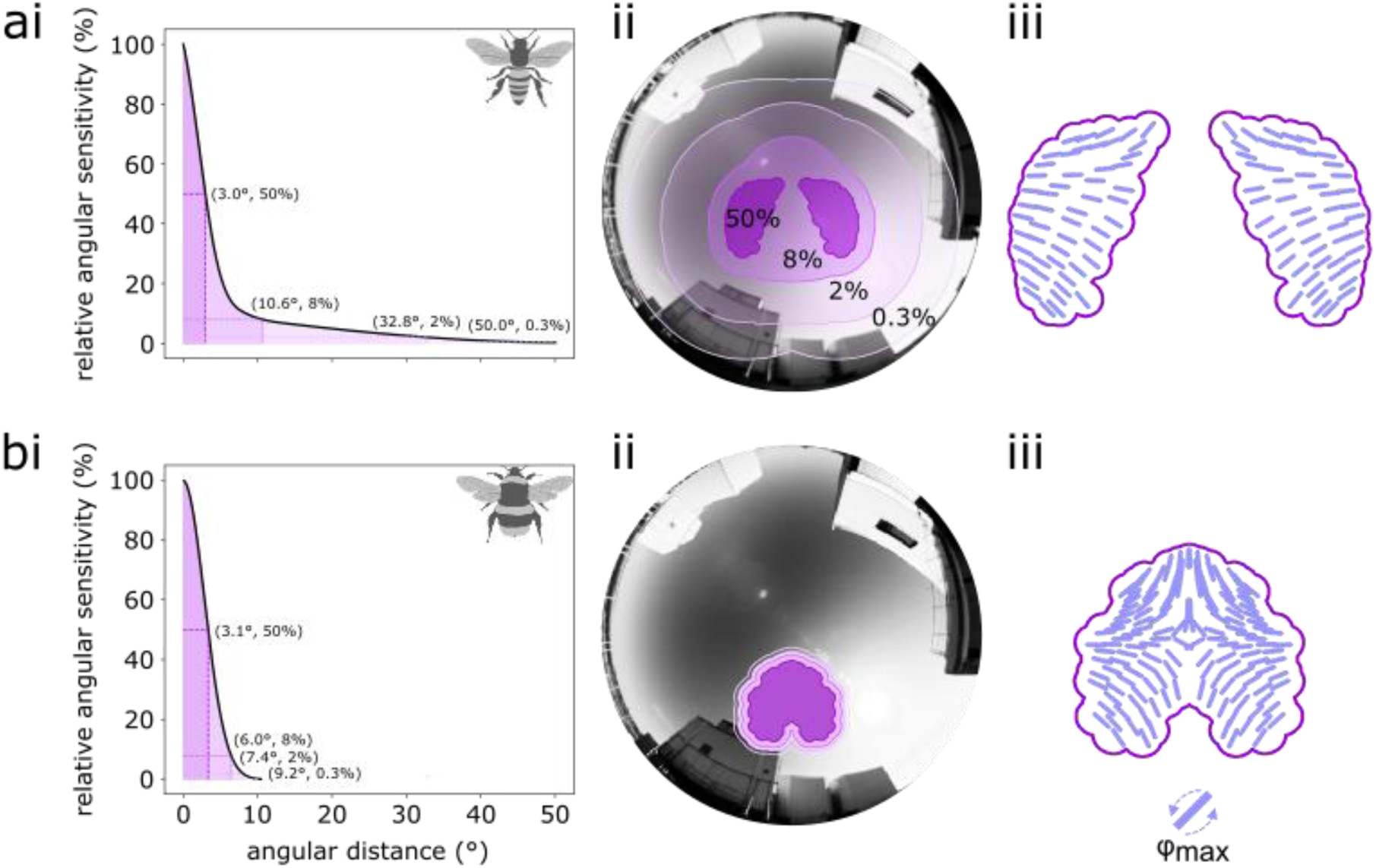
Simulated visual properties of the honeybee (*Apis mellifera*) (a) and bumblebee (*Bombus terrestris*) (b) Dorsal Rim Area (DRA). (i) Angular sensitivity functions of DRA photoreceptors. Both functions were derived from available electrophysiological data (honeybee: Labhart, 1980; bumblebee: Kolyfetis et al., 2025). The honeybee angular sensitivity function (top) is characterised by a wide area of low sensitivity. This low sensitivity area is absent in the bumblebee which is modelled as Gaussian (bottom). Dashed lines correspond to sensitivity levels of 100%-50%, 50%-8%, 8%-2% and 2%-0.3% and the corresponding angular distances are annotated (Half-Width at Half-Maximum). (ii) Total Field of View (FoV) of the DRA. Outlines of the DRA FoV represent different levels of angular sensitivity (corresponding to (i)). The honeybee DRA has a wider FoV than the bumblebee owing to the large low-sensitivity area of individual photoreceptors. (iii) The fan-shaped arrangement (‘filter’) of the photoreceptors’ microvilli is shown for the two eyes of each species. Each ommatidium has one photoreceptor with a preferred AoLP (φ_max1_, shown) to which it is maximally sensitive and a photoreceptor with perpendicular microvilli and an anti-preferred AoLP (φ_max2_, not shown).

Furthermore, we simulated two more, idealised, DRA ommatidial arrangements, namely a dome-shaped one, centred at the zenith^30^ and the same dome shape but tilted by 56° to have minimum overlap with the zenith-looking dome shape. The zenith-centred dome-shaped DRA served as a reference for a DRA with a full 180° azimuth coverage while the forward-facing DRA was used to assess the effect of a forward-facing DRA similar to the bumblebee. For these two ‘idealised DRAs’ we used Gaussian ommatidial FoVs (σ = 2.32°), the same as for the honeybee DRA^22^. For each ommatidium, we calculated a total detected AoLP and DoLP by summing over all pixel intensities of the receptive field and calculating the Stokes parameters.

We assigned a preferred and an anti-preferred AoLP (φ_max1_ and φ_max2_ respectively) to each ommatidium (honeybee: total 106 ommatidia, bumblebee: total 156 ommatidia (Figure 4a_*iii*_,b*_*iii*_*), dome-shaped DRA: total 60 ommatidia). For the honeybee and dome-shaped DRA, φ_max1_ was set to −*az*_*omm*_ + 90° and φ_max2_ was set to −*az*_*omm*_ following Gkanias et al. (in preparation). The bumblebee DRA, which was not centered around the zenith but was more forward-facing (towards the horizon), was first pitched upwards by 29° so that the centre of mass of all ommatidia ([180°,61°]) coincides with the zenith, and then φ_max1_ and φ_max2_ were calculated in the same way. We did this to create a more plausible fan-shaped arrangement of the microvilli which should in theory develop in egocentric coordinates, in the absence of further information. This zenith-based fan-shaped arrangement of the bumblebee would also be anatomically similar to the honeybee one, described in previous studies^22^.

### Matched filter model

To assess the matched filter hypothesis^32,43,44^, we calculated the ‘filter-mismatch’, i.e., the mismatch between AoLP pattern and fan-shaped arrangement (φ_max_ angles; ‘filter’) for our images. For that, we chose the alignment (mismatch) between φ_max2_ and the AoLP pattern (range [0°-90°]) of a ‘composite’ simulated sky image that is comprised by these AoLP that have the maximum DoLP (per pixel) as the sun sweeps from 30°-60° elevation (1° steps; one simulated sky image per step) and 0° sun azimuth (sun behind the animal). We did this since the fan-shaped arrangement is assumed to be a match for the high-DoLP band in the sky for these sun elevations^32,43,44^. φ_max2_ was chosen as the filter for the honeybee DRA as it was calculated to have a better average match with the AoLP pattern in the aforementioned conditions than φ_max1_. More specifically, the honeybee φ_max2_ had an average alignment of 22.4° with the AoLP pattern (77.6° with φ_max1_). For the bumblebee DRA, we also chose alignment with φ_max2_, for consistency, even though its average alignment was 46.48° (43.52° with φ_max1_). The distribution of φ_max2_ angles of our honeybee DRA model (Figure 4a_*iii*_,b*_*iii*_*) also matches well with microvillar axes of R1,5 photoreceptors in the honeybee DRA whereas the distribution of φ_max1_ angles matches well with microvillar axes of R9 photoreceptors^22^. We calculated the arithmetic mean of the filter-mismatch across all ommatidia of each eye and across both eyes, as a measure of deviation from perfect alignment.

We then calculated one response for each of the two photoreceptors with orthogonal axes (R_1_ and R_2_) which were then used to infer one photoreceptor contrast (PRC) for each ommatidium and for the average PRC across both eyes:

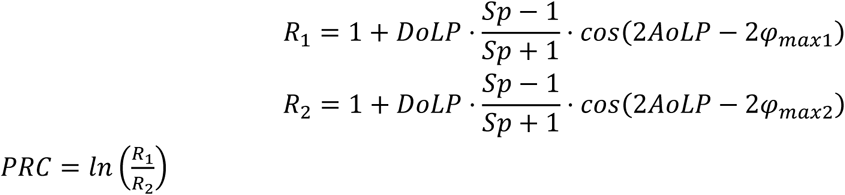

where *Sp* denotes polarisation sensitivity for the DRA photoreceptors^60^. We used *Sp*=6.6 for honeybees^22^ and a slightly higher *Sp*=8 for bumblebees^26^. We used *Sp*=6.3 for the dome-shaped eyes^61^.

To construct heatmaps (Fig. 5-7), images within the same elevation rank (similar sun elevations) were pooled to minimise noise artifacts. The single image at 75° sun elevation was not pooled with any others. Missing values were filled by bicubic interpolation after transformation with an inverse softplus function. The modulation depth across sun azimuths and elevations was calculated as:

**Figure 5.**
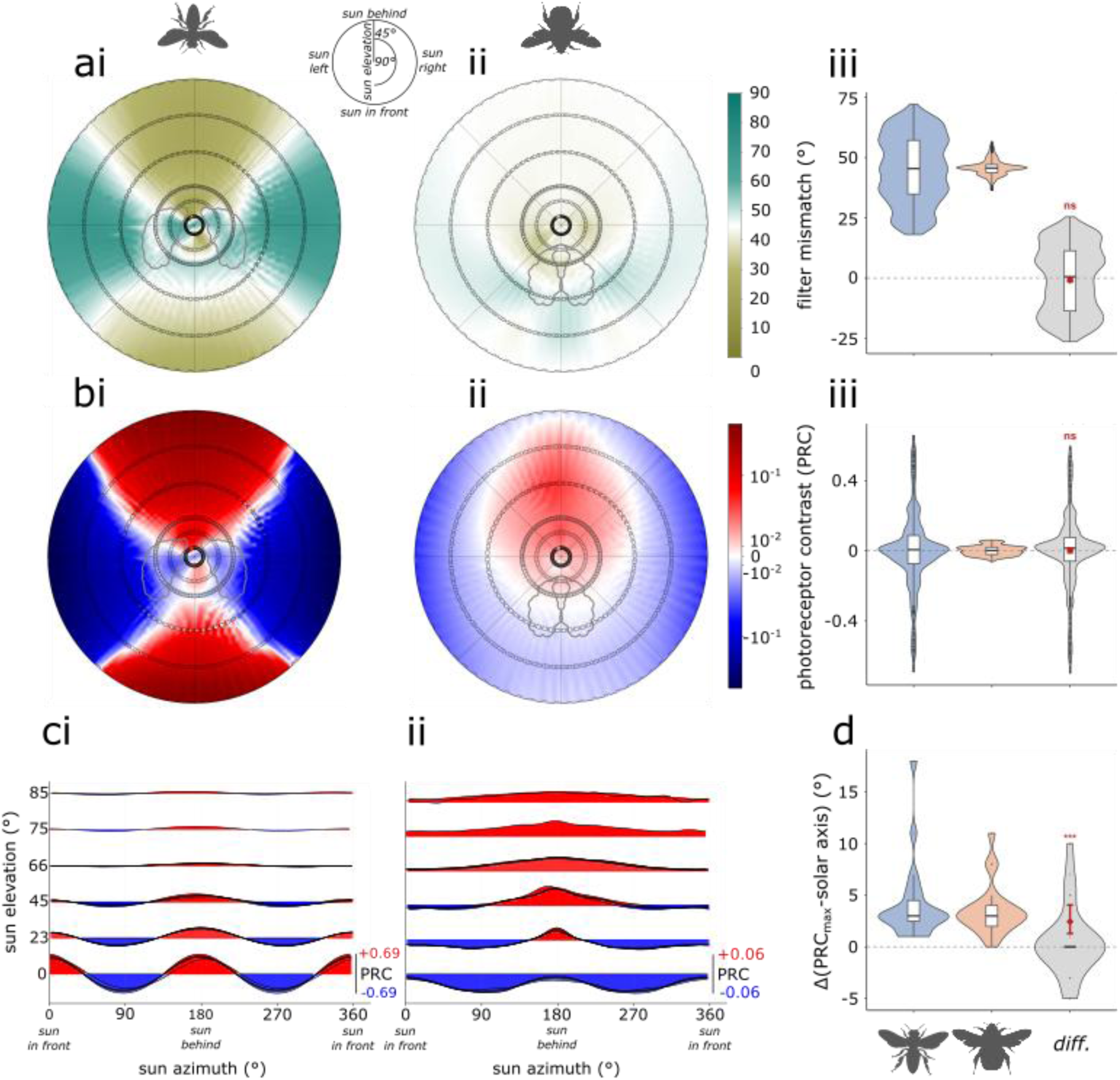
Honeybee and bumblebee matched filter model accuracy. a) Mean filter-mismatch between φ_max_ angles and AoLP for honeybees (i) and bumblebees (ii) across sun positions. (iii) Distribution of filter-mismatch data (not interpolated) for honeybees (left), bumblebees (centre) and their difference (right). b) Mean honeybee (i) and bumblebee (ii) photoreceptor contrast (PRC) across sun positions. (iii) Distribution of PRC data (not interpolated) for honeybees, bumblebees and their difference. c) Ridgeline plots of PRC across sun positions. Honeybee PRC (i) exhibits two peaks when the sun is in front and behind the animal; bumblebee PRC (ii) exhibits a single dominant peak when the sun is behind the animal. Red and blue shading indicates positive and negative PRC, respectively. d) Distribution of the angular offset between the sun position of maximum PRC and the solar axis, for the honeybee, bumblebee and their difference; median angular offset was 3° for both species. The DRA FoV of both species is outlined in grey in the heatmaps. Circles in heatmaps denote data points for six sun elevation ranks (0°, 23°, 45°, 66°, 75° and 85°). For boxplots: red diamond**s** are the model-estimated mean differences (honeybee − bumblebee); red bars are 95% CI; asterisks indicate significance of the overall species effect (Tukey-adjusted: *** p <0.001, ** p <0.01, * p <0.05, ns = not significant).

*modulationdepth* = *max*(*PRC*) − *min*(*PRC*) for each elevation and for 5° azimuth bins (Supplementary Figure 5) as a measure of total response strength.

Accuracy of the matched filter model was assessed by the deviation of the solar axis from the body orientation at which *max*(*PRC*) occurs.

To assess the consistency of the matched-filter model in aligning with the solar axis, we calculated the just-noticeable difference (JND) in rotation from the body orientation of *max*(*PRC*), i.e., how much the animal needs to rotate from *max*(*PRC*) to produce a change in PRC that exceeds the noise. Assuming that bees can follow the general features of the PRC changes across a full body rotation, the PRC across a full body rotation (per-image; 72 body orientations) was smoothed using a Savitzky–Golay filter (window = 11 samples, polynomial order = 3)^62^ as low-pass filter. The body orientation of*max*(*PRC*) was identified from the fitted curve to minimize the effect of outlier values. To define the noise floor, all images of the same elevation rank were pooled and a locally estimated weighted scatterplot smoothing (LOESS) curve was fitted to the pooled data (bandwidth = 30% of pooled data points) with circular padding to ensure continuity at the 0°/360° boundary. The standard deviation of residuals from each elevation-specific LOESS fit was taken as the noise estimate for that elevation rank, reflecting the inter-image variability in PRC that an animal would experience across sky scenes encountered at similar sun elevations. We examined thresholds determined by two possible noise regimes: variable (elevation-specific) and global (across elevation ranks). For the variable noise scenario, the JND for each image was defined as the mean angular distance from the fitted peak required for the PRC to drop by one noise standard deviation, computed by traversing the fitted curve in both directions from the peak and averaging the left and right threshold-crossing distances. For the global noise analysis, the same was done, this time by pooling residuals across all elevation ranks to obtain a single global noise estimate. When the estimated noise exceeded the modulation depth, the threshold fell below the minimum of the fitted curve, resulting in no valid threshold crossing. These cases indicate that signal variation was indistinguishable from noise, and JND was therefore considered undefined and excluded from analysis.

### Vector-sum model

The sun position calculation of the vector-sum model is based on a vector-sum algorithm (Gkanias et al., in preparation), whereby each ommatidium is represented by a vector the direction and magnitude of which are determined by the sign and magnitude of the photoreceptor contrast generated within that ommatidium (positive if aligned more with φ_max1_, negative if aligned more with φ_max2_). Within each ommatidium, vectors associated with negative PRC were oriented 90° relative to those with positive PRC, so that the vector pointed towards the head’s zenith rather than towards the horizon. Then, all vectors from one eye are summed and the resulting vectors from both eyes are themselves summed to a single, total vector that represents the sun azimuth estimate (Figure 3a,b*_i,ii_*,c*_i,ii_*). Note that for the bumblebee for which we used a zenith-based fan-shaped arrangement, the directions of the vectors were based on the zenith-looking ommatidia and not the forward-facing ones. The magnitude of the vectors was determined by the absolute PRC.

To summarise the vector-sum model performance in each image, we sampled each image 72 times across 360° (72 samples, 5° apart), each time acquiring a new sun azimuth estimation. Considering all 72 estimates per image, we calculated a total, average absolute error and a standard deviation for each image, for both species (Figure 3a_*iii*_,b*_*iii*_*).

Moreover, we fitted a mixture of two von Mises distributions to the estimates of each image by optimising a likelihood function (scipy.optimize.minimize; Figure 3a_*iii*_,b*_*iii*_*). We added a weight parameter (p) to the mixture probability density function to distinguish between main and secondary modes:

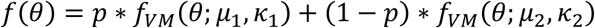

In all conditions, we compared the angular difference of the two modes (μ) of the mixture model to estimate bimodality of the results. Standard deviation (SD) values of the different modes of the mixture von Mises distribution were calculated from the kappa (k) parameter of the modes as follows:

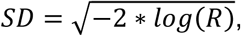

where *R* = *I*1 (*kappa*)⁄*I* 0(*kappa*) and *I*1 and *I*0 are first and zeroth order Bessel functions. For the angular difference of the two modes as well as for the SD values, a spline (scipy.interpolate) was fitted on logit-transformed data. For mode weight parameter values of the mode closest to the sun azimuth (mode_sun_) a spline was fitted to raw values.

### Statistics and visualizations

For performing statistical comparisons of the compass performance between different DRA models (*Apis* vs *Bombus*; dome-shaped DRA vs zenith-centred dome-shaped DRA) and to account for the effects of species, sun elevation, sun azimuth and random effect of images, we fitted linear mixed-effects models (LMMs) using the lme4 package in R, with the exceptions of the JND and solar axis errors of the matched filter model for which ordinary least squares regression was used since each image yielded a single observation (no random effect of images). Post-hoc comparisons between species at each elevation level were performed using the emmeans package with Tukey correction for multiple comparisons. To obtain the overall marginal species effect averaged across elevation levels (predicted contrast; *Apis* - *Bombus*), we extracted estimated marginal means for each species from the best model using emmeans. For more information on model comparisons and post-hoc tests see Supplementary Information (Supplementary Tables 2-16. Supplementary Figures 10-24).

Where undefined JND existed in the JND analysis, the corresponding pairs were also excluded to maintain the same number of paired observations.

All visualizations were done using Python’s Matplotlib^63^, ggplot2^64^ and Inkscape^65^.

## Results

### Sky polarisation imaging

For an accurate account of the skylight polarisation patterns available to foraging bees, we used a dataset of real sky images captured by our polarisation camera and images from a published dataset^53^ across a range of solar elevations. To assess potential differences between real skylight polarisation patterns and predicted simulated ones, we also generated realistic simulated sky dome images^20^, similarly to previous celestial compass models^30^. In terms of physical properties, including AoLP and DoLP, real sky images clearly exhibited greater small-scale variability, despite smoother large-scale gradients for each property (Supplementary Figure 1). Examining disparities between the skylight AoLP patterns for real and simulated skies, it is apparent that these were most pronounced when the sun approached the horizon (where the pixel-wise AoLP deviations reach +/-90°). At higher solar elevations, the patterns were overall more similar, though notable differences remain around the sun and the neutral points^66^, where pattern irregularities are stronger (Supplementary Figure 1).

### Dorsal Rim Area simulation

The anatomical and functional properties of the bee dorsal rim area (DRA) shape the polarisation patterns sampled from the sky and, consequently, the downstream computations underlying polarisation-based navigation. To identify conserved visual features and potential species-specific adaptations, we simulated polarisation vision through the DRA of honeybees (*Apis mellifera*) and bumblebees (*Bombus terrestris*).

The fields of view (FoV) of individual DRA ommatidia were defined by the angular sensitivity function of their photoreceptors (Figure 4a_*i*_,b*_*i*_*) ^22^. The honeybee angular sensitivity function exhibits a sharp peak at the centre of the FoV and a broad area (brim) of low (non-zero) sensitivity around the centre^22^. This is reflected in the large total DRA FoV of the honeybee (Figure 4a_*ii*_). In contrast, the bumblebee angular sensitivity function, modelled as a 2D Gaussian (Figure 4b_*ii*_), does not show this low-sensitivity brim^26^. However, although the total bumblebee DRA FoV is smaller, the Full Width at Half Maximum of the DRA ommatidia receptive fields are similar in the two species, namely 7° for honeybees and 6.24° for bumblebees^26^.

The general viewing direction of the DRA differs between the two species. The honeybee DRA is oriented predominantly upwards toward the zenith (corresponding to 90° sun elevation), whereas the bumblebee DRA is oriented more frontally, toward the animal’s anterior, sampling regions closer to the horizon. The honeybee DRA is composed of fewer ommatidia compared to the bumblebee one (53 and 78 per eye, respectively) but covers a wider range of azimuths (98° and 37° relative to body axis, respectively). In honeybees, the overlap region between the two eyes’ DRAs is confined to low-sensitivity regions (<50%), whereas in bumblebees there is substantially greater binocular overlap, even at high sensitivity levels (>50%). In both species, extensive FoV overlap is also present between neighbouring ommatidia within the same eye, resulting in multiple ommatidia sampling highly overlapping regions of the sky.

The simulated fan-shaped arrangement (‘filter’) of the two species’ DRA photoreceptor microvilli is shown in Figure 4a_*iii*_,b*_*iii*_*. Following our definition for the orientation angle of each ommatidium’s microvilli (φ_max_; see Methods), both species’ fan-shaped arrangements resemble already characterised microvilli arrangements^22,30^ covering a wide range of φ_max_ angles. For honeybees this range was 98° and for bumblebees 177°. The honeybee φ_max_ range roughly matches the already reported range of ∼90°^22^. The bumblebee φ_max_ angle range is wider than the azimuth range (37°) as it was first tilted to the zenith (see Methods).

### Matched filter model accuracy

Bees’ navigational performance depends on accurate estimation of the sun’s azimuth. The matched filter hypothesis proposed by Rossel and Wehner^43^ states that navigational errors of foraging honeybees can be explained by the mismatch between the fan-shaped arrangement of the DRA microvilli (‘filter’) and the AoLP pattern in the sky, a metric termed here as ‘filter-mismatch’. The hypothesis predicts that, as the animal scans the sky, the filter matches best with the AoLP pattern when the animal is aligned with the solar axis (solar-antisolar meridians), i.e., when the sun is directly in front or behind the animal. This alignment also elicits the highest overall ‘response’ or photoreceptor contrast (PRC) of the DRA which allows the animal to accurately track the sun’s azimuth. We attempted to test the main assumptions and implications of this hypothesis for various sun positions using images across six ranks of sun elevation (0°, 23°, 45°, 66°, 75° and 85°).

First, we examined the mean filter-mismatch across sun positions for both species (Figure 5a). We found that, for honeybees, the filter-mismatch (Figure 5a_*i*_) is indeed low when the sun is located on the anterior–posterior axis, largely independent of solar elevation, in agreement with the matched filter hypothesis. The filter-mismatch becomes higher when the sun is located on the lateral axis of the animal, again largely independent of solar elevation. Note that the filter-mismatch is measured from φ_max2_ angles so that a high filter-mismatch is low with φ_max1_ angles since the hypothesis implies that it is a single class of photoreceptors that has to match the AoLP pattern in the sky^43,44^. The bumblebee pattern of filter-mismatch across sun positions (Figure 5a_*ii*_) differs from the honeybee one. Filter-mismatch values are generally intermediate (≈45°), especially when the sun is behind the animal (0°-45° sun elevations). This is likely a result of the higher range of φ_max_ angles within each eye (Figure 4b_*iii*_) which lead some parts of the eye to be best aligned with one φ_max_ angle and others with its perpendicular. Higher values of filter-mismatch are observed when the sun is in front of the animal at elevations of 45°-85° while the values are lower for sun slightly behind the animal. Both species have filter-mismatch values evenly distributed around 45° (honeybees: median=45.4°; bumblebees: median=45.6°; predicted contrast=-0.7°) with bumblebees having a narrower range of values compared to honeybees (honeybees: IQR=22.2°; bumblebees: IQR= 3.3°) (Figure 5a_*iii*_), again as a result of the narrow φ_max_ angle range.

We also examined the summed ‘response’ or photoreceptor contrast (PRC) of the DRA across sun positions (Figure 5b). According to the hypothesis, PRC ought to be highest when the animal is aligned with the solar axis, along with corresponding low filter-mismatch^43,44^. In agreement with the hypothesis, we see that, for honeybees, PRC is strongly correlated with filter-mismatch (Figure 5b_*i*_). We see strong, positive overall responses when the sun is in front or behind the animal, independent of sun elevation. Conversely, we see strong negative overall responses when the sun is to the left or right of the animal. The switch from positive to negative PRC values is sharp and occurs when the sun is roughly located obliquely to the animal’s body axis. Responses are overall weaker (around 10-fold) for bumblebees compared to honeybees, which is consistent with weaker overall alignment of the filter with the AoLP pattern. Bumblebees’ PRC values (Figure 5b_*ii*_) are mostly positive when the sun is behind the animal, reaching a maximum at 45° elevation. Responses are mostly negative for all other sun positions. For both species, the absolute PRC values are generally low for elevations greater than 45° or when the sun overlaps with the FoV of the DRA, owing to the low DoLP around the sun. Both species have PRC values evenly distributed around 0 (honeybees: median=0.007; bumblebees: median=0.002; predicted contrast=0.003) with bumblebees having a narrower range of values compared to honeybees (honeybees: IQR=0.161; bumblebees: IQR= 0.04) (Figure 5b_*iii*_).

To assess the matched filter in terms of accuracy, we examined the angular deviation between the azimuth that corresponds to maximum PRC and the solar axis. PRC change as a function of sun position can be seen in Figure 5c. In agreement with the matched filter hypothesis, the maximum PRC occurs, for both species, when the animal is aligned either with the sun (0°; sun in front) or the anti-sun (180°; sun behind) (Figure 5c_*i*_*_,ii_*). The overall amplitude of the PRC decreases as sun elevation increases, likely because of decreasing DoLP. However, the shape of the response function does not change with sun elevation. For honeybees (Figure 5c_*i*_), the shape is sinusoidal with peaks at 0° and 180° and troughs at 90° and 270°. As a result of the sinusoidal shape, each PRC value (except for the maximum and minimum) occurs at least two times while the animal scans the AoLP pattern. Thus, orientation at most angles relative to the sun is ambiguous. The bumblebee PRC function is only sinusoidal at low sun elevations (0°; Figure 5c_*ii*_). In all other cases, PRC increases almost linearly as the sun moves from the front to the back of the animal. The single peak is likely a result of the two eyes viewing very similar parts of the sky in the bumblebee which is not the case in the honeybee. The median angular deviation of maximum PRC from the solar axis is 3° for both species and the predicted contrast is 2.44° (Figure 5d). Interestingly, maximum PRC occurs when the sun is behind the animal in most cases, for both species. This is the case because the eyes of both species overlap with the high DoLP region in the sky when the sun is behind them. There are five exceptions to that trend (3 for honeybees and 2 for bumblebees), when the sun is at the horizon, where maximum PRC occurs when the sun is in front of the animal. This shows that when the sun is at the horizon, solar axis ambiguity increases due to the high symmetry in the AoLP pattern at that time. Overall, the matched filter model succeeds in identifying the solar axis with high accuracy in both species, even though PRC across sun positions differs notably between species both in amplitude and distribution.

Identical analyses on simulated skies are broadly similar at intermediate sun elevations but reveal distinct differences compared to real skies, especially at low sun elevations (see Supplementary Figure 6).

Furthermore, we calculated the modulation depths for both species, both across sun azimuths and elevations (Supplementary Figure 5). The modulation depth across azimuths indicates the level of change in PRC as the animal changes its body orientation whereas across elevations it shows how sensitive PRC is to sun elevation. Across azimuths (5° bins), for a given elevation, honeybee modulation depths exhibit four maxima at 0°, 90°, 180° and 270°, which coincide with the highest (positive) and lowest (negative) PRC values. There are four minima at 45°, 135°, 225° and 315° which coincide with intermediate (≈0) PRC values. There, the two photoreceptor classes have approximately equal responses, or the responses cancel out across ommatidia. Across sun elevations, the modulation depth decreases steadily from low to high elevations, indicating that differences in PRC are more subtle at high sun elevations.

For bumblebees, there are only two maxima at 90° and 270° and one minimum at 180° sun azimuth which again coincide with high/low and more uniform (positive) PRC values, respectively. Across sun elevations, modulation depths exhibit a peak at ≈45° sun elevation which coincides with a local increase of PRC values when the sun is behind the animal and a decrease of PRC values when the sun is in front, possibly corresponding to the animal viewing the maximum DoLP band in the sky. Modulation depth values are decreased near the zenith where PRC is mainly positive (Supplementary Figure 5).

### Vector-sum model accuracy

We also examined an alternative sun azimuth estimation mechanism, the vector-sum model (Gkanias et al., in preparation). Unlike the matched filter model, which requires the animal to scan the sky in multiple body orientations to identify the solar axis, the vector-sum model produces an instantaneous sun azimuth estimate from a single body orientation by summing PRC values as vectors.

For honeybees, absolute angular errors of the vector-sum model increase (>90°) as the sun approaches the horizon (elevations 0°–23°) from the front or from posterior left and right of the animal (Figure 6a_*i*_). The errors are also high (≈90°) when the sun approaches the zenith (directly above the animal). When the sun is positioned in all other regions of the sky, the errors are relatively low (<30°). For bumblebees (Figure 6a_*ii*_), errors are generally high (>90°) when the sun is located behind the animal, especially at low elevations (<23°). Errors are also high when the sun overlaps with the DRA FoV (directly above the animal), especially in the overlap region of the two eyes, and when it is near the zenith. When the sun is located in front of the animal, errors are low (<30°). Across all sun positions, honeybees exhibit a lower median error compared to bumblebees (honeybees: median=10.8°; bumblebees: median=28.2°; predicted contrast=-25.01°) (Figure 6a_*iii*_). It is worth noting that the high error (∼90°) regions in Figure 5a may be a result of interpolation between datapoints on the heatmap, as analyses on simulated skies suggest (Supplementary Figure 7).

**Figure 6.**
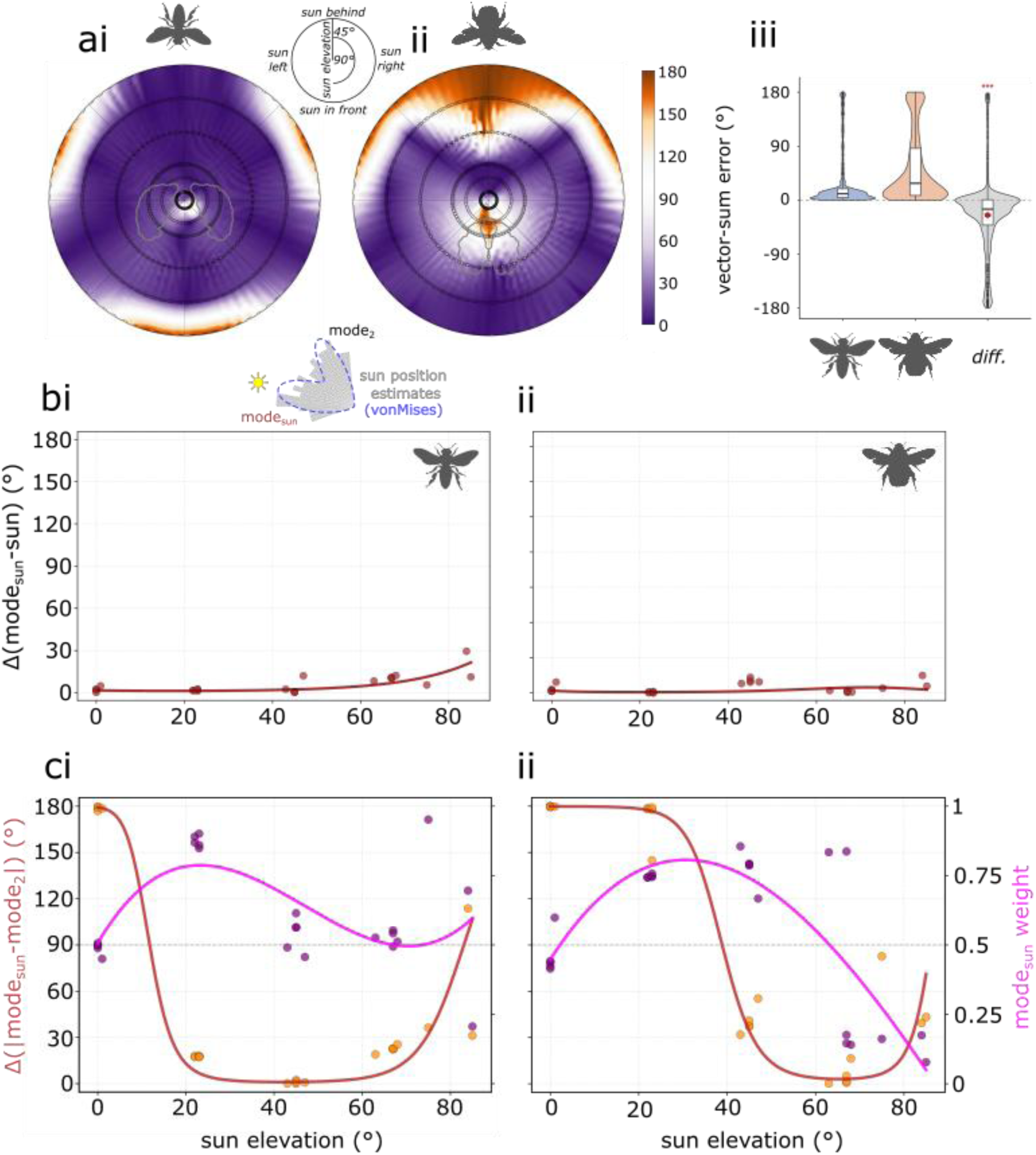
Honeybee and bumblebee vector-sum model accuracy. a) Mean absolute error of the vector-sum model for honeybees (i) and bumblebees (ii) across sun positions. (iii) Distribution of errors (not interpolated) for honeybees (left), bumblebees (centre) and their difference (right). The DRA FoV of each species is outlined in grey in the heatmaps. Circles in heatmaps denote data points for six sun elevation ranks (0°, 23°, 45°, 66°, 75° and 85°). Differences are honeybee minus bumblebee. b) Angular deviation of the mixture von Mises mode closest to the sun azimuth from the true sun azimuth, as a function of sun elevation, for honeybees (i) and bumblebees (ii). c) Bimodality of vector-sum estimates shown as the absolute angular difference between the two modes of the mixture von Mises fit (left axis, red) and the weighting parameter (*p*) of the mode closest to the sun azimuth (right axis, magenta), as a function of sun elevation, for honeybees (i) and bumblebees (ii). Lines show spline fits to logit-transformed values (b-c; red) or raw values (c; magenta). For boxplots: red diamond**s** are the model-estimated mean differences (honeybee − bumblebee); red bars are 95% CI; asterisks indicate significance of the overall species effect (Tukey-adjusted: *** p <0.001, ** p <0.01, * p <0.05, ns = not significant).

**Figure 7.**
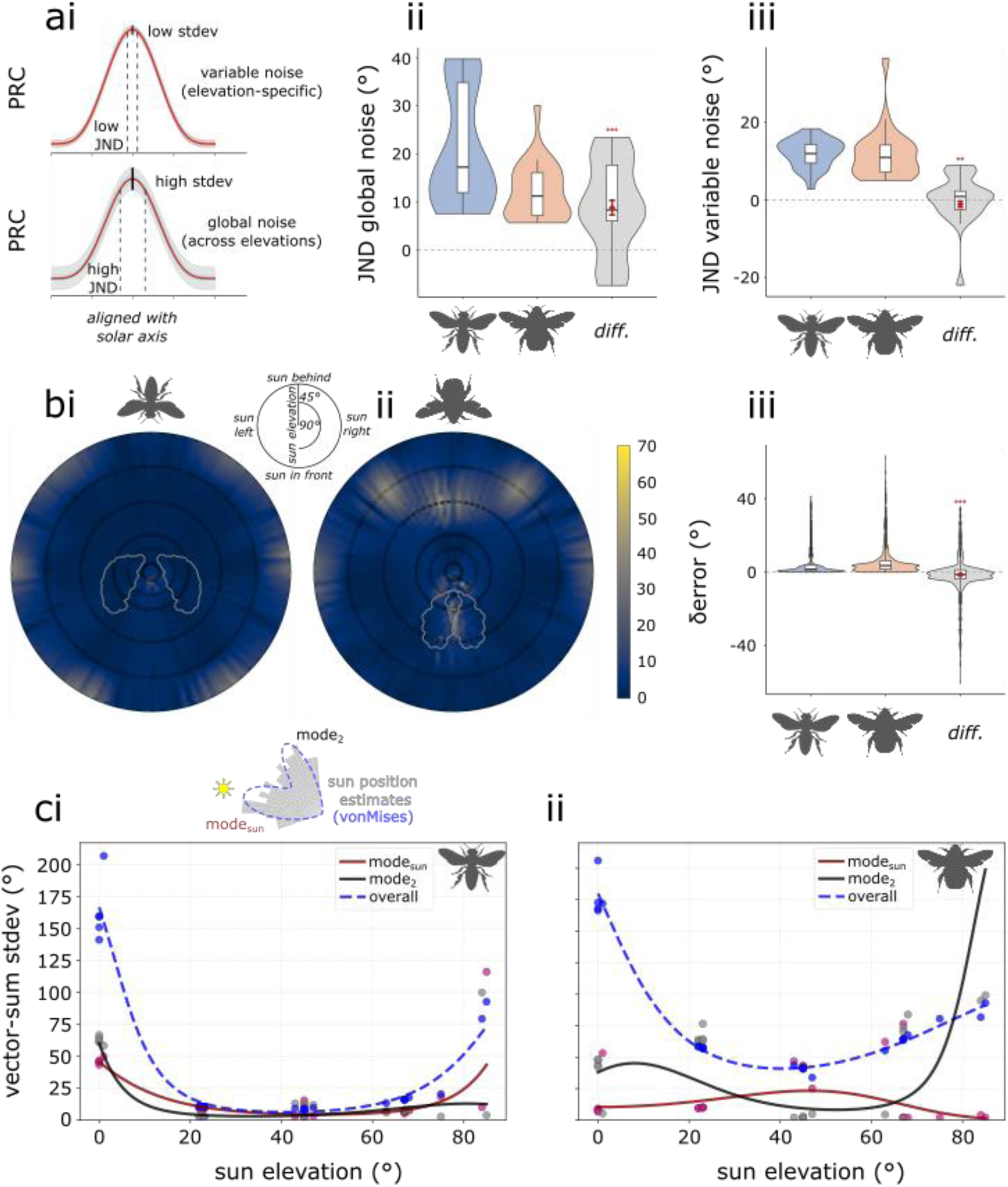
Course control stability of the matched filter (a) and the vector-sum (b,c) models. (a) Schematic illustrating the just-noticeable difference (JND) in body rotation from maximum photoreceptor contrast (PRC) under variable, elevation-specific noise (top) and global noise across all elevations (bottom). A higher standard deviation (SD) of PRC values results in a higher JND. (ii) Distribution of JND values under global noise for honeybees (left), bumblebees (center), and their difference (right). (iii) Distribution of JND values under variable noise for honeybees, bumblebees, and their difference. (b) Mean absolute error difference (δerror) between adjacent sun positions (5° apart in azimuth) for honeybees (i) and bumblebees (ii) across sun positions. (iii) Distribution of δerror values for honeybees, bumblebees, and their difference. (c) Standard deviation of vector-sum sun position estimates as a function of sun elevation for honeybees (i) and bumblebees (ii), shown separately for the mode closest to the sun azimuth (mode_sun_; red), the other mode (mode_2_; black), and the total estimate distribution (overall; dashed blue). Points represent individual images. The DRA of both species is outlined in grey in the heatmaps. Circles in heatmaps denote uninterpolated data points at six sun elevation ranks (0°, 23°, 45°, 66°, 75°, 85°). For boxplots: red diamond**s** are the model-estimated mean differences (honeybee − bumblebee); red bars are 95% CI; asterisks indicate significance of the overall species effect (Tukey-adjusted: *** p <0.001, ** p <0.01, * p <0.05, ns = not significant).

To summarise the model’s performance at each sun elevation, we fitted a two-component von Mises mixture distribution to sun azimuth estimates across body orientations. Across both species and most elevations, deviations between the true sun azimuth and the nearest mode mean (mode_sun_) were typically <10°, indicating good performance (Figure 6b). Performance decreased at high sun elevations for honeybees, with deviations reaching ∼25° at 85° (Figure 6b_*i*_). Bumblebees’ performance decreased slightly at 45° and 85° with mode_sun_ deviations of ∼20°. Notably, low mode_sun_ deviations often coincided with high absolute errors (Figure 6a). For example, honeybees showed large absolute errors at low sun elevations, and bumblebees when the sun was posterior, yet these effects were not reflected in the mode_sun_ deviations (Figure 6b). This discrepancy arises because the mean absolute error averages across all estimates for a given image, whereas the mode_sun_ deviation reflects only those estimates clustering around the true sun azimuth. This reflects that increased vector-sum errors may be the result of ambiguity between two opposing directions.

Accurately estimating the solar axis is a crucial feature of the polarisation compass, since the exact sun azimuth can then be recovered using complementary visual cues. Given that one mode (mode_sun_) closely tracks the true sun azimuth (Figure 6b), we quantified axial error (solar axis deviation) as the angular separation between the two modes (mode_sun_ and mode_2_) of the fitted von Mises mixture (Figure 6c; red). Large mode separations imply that the estimate lies far from the solar axis, yielding increased axial error, whereas small separations indicate low axial error. Honeybees exhibit high axial error (>20° mode separation) when the sun is near the horizon or approaches the zenith, with low axial error at intermediate elevations (Figure 6c_*i*_). Bumblebees show very high axial error (∼180° mode separation) at low sun elevations (<45°), which decreases sharply above 45° and reaches low values at higher elevations (Figure 6c_*ii*_). Axial errors across azimuths for both species can also be seen in Figure 6a_*i*_*_,ii_* where white regions are associated with absolute errors near 90°, i.e., high axial error.

To investigate whether the high axial errors are an effect of the forward-facing DRA, we conducted the same analyses on the idealised, forward facing, dome-shaped DRA. The trend is similar to the bumblebee with the exception that axial error is increased at high sun elevations (Supplementary Figure 8).

To further characterize how compass estimates are distributed between competing directions, we examined the weighting parameter of the von Mises mixture model. The weighting parameter (*p*_*sun*_; range 0–1) describes the contribution of the mode closest to the sun azimuth (mode_sun_) to the mixture, corresponding to the probability that an estimate clusters around that mode rather than the competing direction (Figure 6c; magenta). In honeybees, *p*_*sun*_ remains near 0.5 at most sun elevations, indicating no strong bias towards either mode, except at ∼23° elevation where mode_sun_ is favoured. Importantly, strong bimodality occurs only when the sun is near the horizon or close to the zenith, consistent with the high axial errors observed at these elevations (Figure 6c). In bumblebees, *p*_*sun*_ is ∼0.5 near the horizon; it increases across intermediate elevations (23°-66°), where axial error is generally high, and decreases again as the sun approaches the zenith, where axial error is low.

Similarly to the matched filter model, identical analyses on simulated skies are consistent with real skies for intermediate elevations, but differences arise at low elevations for both species (Supplementary Figure 7).

### Course control stability

Successful navigation requires not only accurate estimation of the sun azimuth, but also the ability to maintain a stable course while moving by keeping the sun (or solar axis) at a constant angle relative to the body axis.

To quantify polarisation-based course control stability for the matched filter model, we measured the just-noticeable difference (JND) in body rotation from maximum PRC that can elicit a change in PRC greater than the noise (residual standard deviation (SD) of PRC values). Generally, a larger JND indicates lower sensitivity to changes in body rotation, so that larger angular deviations are required before differences in PRC exceed the noise threshold, leading to less precise course adjustments. Conversely, a smaller JND reflects higher sensitivity, enabling finer corrections, and thus greater course control accuracy.

We examined both global (across all elevations) and variable (elevation-specific) noise (Figure 7a_*i*_). With the assumption of global noise, JND is generally larger as a result of greater SD of PRC values (Figure 7a_*ii*_). Honeybees exhibit higher JND across sun elevations than bumblebees (honeybees: median=17.3°; bumblebees: median=11.3°; predicted contrast=8.8°) and a wider range of JND values (honeybees: IQR=22.8°; bumblebees: IQR=8.9°). The higher JND values of honeybees likely result from the wide range of PRC values across elevations (Figure 5b,c) which lead to increased SD. Bumblebees’ lower JND likely reflects their narrower range of PRC values (Figure 5b,c), resulting in lower SD. With the assumption of variable, elevation-specific noise, both species have similar JND values (honeybees: median=11.9°; bumblebees: median=10.9°; predicted contrast=-1.2°) and ranges (honeybees: IQR=4.8°; bumblebees: IQR=7°). Thus, assuming that no additional noise is added to course control when the animal rotates away from the aligned orientation, both species can maintain stable course with deviations under 20°.

We quantified course control stability for the vector-sum model in two ways: the absolute error difference (δerror) between adjacent sun positions (5° apart in azimuth; Figure 7b) and the SD of sun position estimates per image (Figure 7c). δerror indicates how a small change in body orientation affects the sun position estimate and thus the ability to maintain a stable course while the standard deviation of estimates summarises the consistency of the model for a given sun elevation. δerror is generally low (<5°) across sun positions (Figure 7b_*i*_*_,ii_*) for both species (honeybees: median=1.6°; bumblebees: median=3.6°; predicted contrast=-1.36°; Figure 7b_*iii*_). Some exceptions arise for the honeybees at low sun elevations (δerror≈40°) and for bumblebees at intermediate sun elevations (23°-45°; δerror≈45°) when the sun is behind the animal. δerror values suggest that both species can maintain a stable course across most sun positions, with the largest instabilities occurring at the same elevations where absolute error is high (Figure 6a).

The SD of sun azimuth estimates was measured, per image, for both von Mises modes (mode_sun_ and mode_2_) separately and for the total estimates (overall; Figure 7c). Both species exhibit highly consistent estimates under intermediate sun elevations, as indicated by low SD values. In honeybees (Figure 7c_*i*_), SD is low (<25°) between ∼23° and 75° sun elevation but increases at very low (0°) and very high (>75°) elevations, indicating reduced consistency near sunrise/sunset and when the sun approaches the zenith. Bumblebees show a gradual decrease in SD with increasing sun elevation, stabilising at a SD of ∼75° for elevations above 45°, which suggests improved consistency as the sun rises (Figure 7c_*ii*_). High SD values for low sun elevations are likely the result of the estimates being bimodally distributed around the sun and anti-sun. In both species, SD values computed for individual von Mises modes are consistently lower than those for the full estimate distributions. Overall, the vector-sum model produces consistent estimates for both species at intermediate sun elevations, with individual modes remaining stable even when the total estimate distribution is broad; consistency is reduced at low and high elevations where absolute error is also highest. These results are comparable with results from simulated skies, albeit with distinct differences at low sun elevations (Supplementary Figure 9).

## Discussion

### A biologically accurate model of the bee DRA & polarisation-based navigation

Many insects, including bees, perceive polarised skylight through the dorsal rim area (DRA) of their compound eyes to estimate the sun’s azimuth, which allows them to navigate accurately. Here, we developed a biologically grounded simulation of polarisation vision in honeybees and bumblebees, incorporating species-specific DRA anatomy and real sky polarisation images. We evaluated two alternative, not mutually exclusive, sun azimuth estimation mechanisms: the matched filter and vector-sum models. Although both models concern orientation based on sun azimuth estimation, the matched filter produces only one estimate per sun position, i.e., per image, as it requires scanning of the whole sky, whereas the vector-sum model can produce multiple estimates per sun position, depending on body orientation. Thus, the two models allow only for similar, but not identical measures of performance.

The main findings of this study can be summarised as follows. Real sky images reveal that simulated skies overestimate available polarisation information when the sun is near the horizon. Species-specific differences in DRA viewing direction constitute the main driver of navigational performance differences between honeybees and bumblebees. While the matched filter accurately identifies body alignment mainly with the anti-solar meridian in both species, it requires active scanning across body orientations to do so, a fundamental limitation compared to the vector-sum model, which generates instantaneous sun position estimates from any single body orientation. Both models allow stable course control in both species and under most sun elevations. However, the matched filter being limited to solar axis alignment, only enables a form of consistent, positive or negative phototaxis. Together, these results offer a mechanistic and comparative framework linking DRA anatomy to navigational performance under natural sky conditions.

### Polarisation sky images

Insects that rely on the sun to navigate use multiple skylight cues to determine its position including brightness and spectral gradients and the polarisation pattern (e.g. ^67^). Previous studies, aimed at modelling the insect solar compass, used simulated sky conditions (e.g. ^30^) that assume high uniformity across the sky dome with respect to the spatial structure of light properties. For instance, the Rayleigh sky model predicts a structured, radially symmetric, sun-centred polarisation pattern^18^. However, empirical measurements show that real skylight polarisation patterns deviate from this theoretical model. More realistic sky models incorporate additional atmospheric effects (e.g., multiple scattering and inhomogeneities), and produce polarisation patterns that are nonlinear with respect to both angular distance from and direction to the sun, and are thus less radially symmetric^19^. These deviations result in irregularities in both DoLP and AoLP that are not captured by the Rayleigh model but are more characteristic of natural skies. Nevertheless, even these more realistic simulations make assumptions of spatial uniformity that may not reflect natural skies.

The need for assumptions about the spatial organisation of sky polarisation patterns can be removed by leveraging real sky images, something that has rarely been done (e.g., ^50^). Our results reveal considerable differences between real and simulated skylight AoLP patterns, particularly at low sun elevations; times of high ecological relevance, as both honeybees and bumblebees forage from early morning until late evening^68,69^. AoLP in real skies becomes roughly parallel when the sun is near the horizon, in contrast to the modelled radially symmetric structure of the simulated skies. Thus, simulated patterns likely overestimate the available polarisation information at low sun elevations (Supplementary Figure 1). Indeed, supplementary analyses confirm that it is at low sun elevations that we observe the most notable differences in both species’ matched filter and vector-sum model performance between real and simulated skies (Figures 5,6; Supplementary Figures 6,7,9). These findings suggest that simulated skies provide a reasonable approximation at intermediate sun elevations, but that real sky images may be necessary to model insect navigation accurately during all ecologically relevant periods including near sunrise and sunset.

### DRA visual properties

#### Angular sensitivity

The angular sensitivity or acceptance function of the DRA ommatidia, i.e., the relative probability of an ommatidium capturing a photon at each angular distance from the centre of the FoV of the ommatidium, is usually approximated by a radially symmetric Gaussian function^70,71^. However, electrophysiological studies suggest that this may be inappropriate for honeybees^22^. Here, we used an accurate the acceptance function of honeybee DRA photoreceptors derived from electrophysiological data^22^ (Figure 4a_*i*_). For the bumblebee DRA ommatidia, absent similar data for angular sensitivities at large off-axis angles, we used Gaussian acceptance functions^26^. The absence of pore canals in the corneal lenses of *Bombus terrestris* DRA is consistent with the absence of a wide brim in the FoV, as pore canals are thought to underlie the brim observed in honeybees^22^.

The angular sensitivity curve of honeybee DRA photoreceptors shows a high relative sensitivity at the centre of the ommatidial FoV and a wide, low sensitivity area (brim; ∼30° radius) around the center^22^. The vector-sum model performance was, however, similar to that of a DRA composed of ommatidia with Gaussian receptive fields (Supplementary Figure 4). This result suggests that a Gaussian receptive field suffices to provide the necessary polarisation information to a single ommatidium at least under clear sky conditions, and a wider receptive field might offer diminishing returns in compass performance. The benefits of wide receptive fields, which may include the smoothing of noise present in the polarisation pattern, are expected to be greatest under cloudy conditions^72^, although this effect remains to be investigated in greater detail in future studies.

#### DRA viewing direction

The general viewing direction of the DRA plays a crucial role in determining which part of the sky bees sample from while flying. Here, we used the surface normal projections of the honeybee DRA facets as a proxy for the centre of each facet’s viewing direction. The DRA viewing surface inferred in this way matched already reported data on the FoV of the honeybee DRA^22^ (Supplementary Figure 2). Honeybees and bumblebees differ notably in the viewing direction of their DRA. The honeybee DRA is oriented more towards the zenith compared to that of the bumblebee, which has a more frontal FoV (tilted towards the horizon). This influences the time of day at which the DRA’s FoV overlaps with the sun while flying. For example, a DRA’s FoV centred at 30° elevation will overlap with the sun mostly when the sun is also at 30° elevation. The time of day when that happens may vary between different times of year and localities but is consistent for a given locality. Angles of polarisation change more dramatically throughout the day in a sky patch near the zenith compared to a sky patch near the horizon, where angles change more gradually. Consequently, a DRA viewing the zenith (like the honeybee DRA) observes a wider range of AoLP than one with a lower FoV (like the bumblebee DRA).

A zenith-looking DRA may reflect an effective adaptation that maximises view of high-DoLP regions of the sky for the solar elevations most encountered by the two species. Honeybees (*Apis mellifera*) have a geographical distribution that includes tropical and temperate regions while bumblebees (*Bombus terrestris*) are distributed in temperate and sub-arctic regions^73^. This means that they mostly forage in regions where the sun never reaches elevations above ∼75° (temperate and sub-arctic regions). Even when it does reach such high elevations (tropical regions), it spends relatively less time near the zenith compared to lower elevations. As a result, a zenith-looking DRA is generally less likely to view the bright region near the sun where DoLP, i.e., the signal, are typically much lower. Importantly, DoLP is critical for accurate recovery of the AoLP^72^, which carries the primary information on solar azimuth, and a major driver of photoreceptor contrast^60^ which in turn is the input for any solar compass model.

The two advantages of a zenith-oriented DRA, i.e., sampling of a wider AoLP range and reduced overlap with low-DoLP regions near the sun, are consistent with the overall lower absolute errors of the honeybee vector-sum model and the higher errors of the bumblebee (Figure 6). They are also consistent with the higher errors of the forward-facing dome-shaped DRA compared to the zenith-facing one (Supplementary Figure 8). A tilted, dome-shaped DRA has been shown to lead to decreased sun position estimation accuracy^30^. In this study, we observe the same effect of tilt in the bumblebee DRA (Figure 6). Whether DRA viewing direction and anatomy more broadly correlates with species-specific foraging patterns, such as time of day and latitude of activity, remains an open question. A systematic comparative investigation coupling DRA anatomy with detailed behavioural observations could reveal the ecological drivers and evolutionary pressures shaping ocular anatomy across species.

### Revisiting the matched filter hypothesis

Microvillar orientation within the DRA is a further anatomical feature that can affect insect polarisation vision and orientation. The fan-shaped microvillar arrangement of the ant and honeybee DRA ommatidia has been hypothesised to match the average AoLP pattern in the sky^32,43,44^, acting as a built-in matched filter in the ‘hardware’ of the compound eye. According to the matched filter hypothesis, by rotating their bodies animals can scan the sky dome, identifying the body orientation at which the fan-shaped arrangement of the microvilli best matches the AoLP pattern, resulting in the maximum overall DRA photoreceptor response^44^. The maximum match is hypothesised to be achieved when the animal is aligned with the symmetry plane of the skylight AoLP pattern, i.e., with the solar axis^44,74^. This scanning behaviour constitutes a form of active vision, a concept that has been studied in a variety of visual tasks in bees^75–78^. The matched filter mechanism relies on actively manipulating the PRC output and achieves a reliable and steady alignment with the solar axis, something that cannot be achieved through passive vision under this model.

#### Quantification

Our simulation allowed us to quantify the average match between the fan-shaped microvillar arrangement (‘filter’) of the DRA and the skylight AoLP pattern, here called filter-mismatch. This filter-mismatch is hypothesised to be low only for a small part of the DRA in honeybees^79^, crickets^80^ and locusts^33^.

As Figure 5a_*i*_ suggests, the lowest overall filter-mismatches for our honeybee DRA model are indeed achieved when the sun is located on the anterior-posterior axis, largely independent of sun elevation, in agreement with the matched filter hypothesis^43^. The honeybee PRC pattern across sun azimuths and elevations (Figure 5b_*i*_) strongly resembles this filter-mismatch pattern. Since PRC is driven by both filter-mismatch and DoLP (see Methods), this suggests that filter-mismatch dominates PRC calculation and minimises the effect of DoLP differences across ommatidia and eyes.

For the bumblebee DRA model, however, there is not a strong correspondence between the overall filter-mismatch (Figure 5a_*ii*_) and the PRC pattern (Figure 5b_*ii*_). The generally intermediate mismatch (∼45°) is probably a result of the wide range of φ_max_ angles which may neutralize the effect of the filter-mismatch in the calculation of the PRC. The higher positive PRC observed when the sun is behind the animal in both species (Figure 5c) is consistent with the DRA’s frontal tilt (particularly pronounced in bumblebees) which maximises overlap between the DRA FoV and the high-DoLP band in the sky at those sun positions.

#### Accuracy

The accuracy of the matched filter in determining the solar axis can be assessed by the amplitude of PRC across sun positions and its alignment with the solar axis. If we assume that POL-neurons exhibit a constant noise threshold independent of sun elevation, i.e. that their ability to detect maximum PRC depends on the PRC amplitude, then honeybees would face the most difficulties in aligning with the solar axis at high sun elevations where PRC amplitude is lowest (due to increased FoV overlap with the sun) even though the maximum PRC still aligns with the solar axis (Figure 5c_*i*_). On the contrary, bumblebees show smaller amplitude differences in PRC across elevations, even though PRC is generally lower than the honeybee. It can be predicted then that they would align equally well with the solar axis at low and high sun elevations. If we assume that POL-neurons can reliably detect maximum PRC when scanning the sky, i.e., independent of amplitude, then both species can very accurately align with the solar axis, and predominantly with the anti-solar meridian, with less than 5° error (Figure 5d). Resolving the ambiguity between solar and anti-solar alignment (when peak PRC values are close) could be achieved through the use of complementary cues, such as spectral information^44^.

It is worth noting that the shape of the PRC distribution across body orientations for a given sun elevation differs between the two species (Figure 5c) as a result of DRA anatomy. Honeybees’ PRC is sinusoidal across azimuths with two peaks of similar amplitudes when the sun is in front and behind the animal, because of the greater symmetry of the DRA along both left-right and anterior-posterior axes. Conversely, the bumblebee DRA produces a single peak when the sun is behind (but not at the horizon). This presents a potential advantage of the forward-facing bumblebee DRA since the DoLP difference between sun in front and behind is greater and aids in disambiguating the alignment. Another anatomical characteristic that factors in this disambiguation is the fact that the DRA of the two bumblebee eyes sample similar sky regions with high binocular overlap, producing similar PRC values, which is not the case for the honeybee. Overall, differences in DRA anatomy lead to distinct PRC patterns across sun positions that may influence the alignment accuracy with the solar axis and potentially the ability to resolve solar versus anti-solar meridians.

#### Consistency

Our results show that the matched filter can be used by bees to maintain a stable course while flying, a crucial navigational task for many insects^81,82^. To achieve this, bees would have to first align with the solar axis and then make sure that they do not deviate from that alignment while flying, a form of phototaxis^5^. We have quantified this by calculating the just-noticeable difference (JND) in body rotation based on the PRC across sun positions, i.e., the minimum rotation needed for a detectable (larger than the noise level) difference in PRC. By considering two possible models for inter-image noise (SD of PRC), we account for the fact that hypothetical ‘rotation detector neurons’ may be tuned to either noise across all possible sun elevations (global) or to elevation-specific (variable) noise.

JND values are higher under global noise for both species compared to variable noise (Figure 7a) since the standard deviation of PRC under global noise is calculated across images of all sun elevations. Interestingly, the difference is more pronounced in honeybees (median_global_=17.3° vs median_variable_=11.9°) while bumblebees exhibit similar JND values under both noise models (median_global_=11.3° vs median_variable_=10.9°). This presents another advantage of the bumblebee DRA because PRC values are more consistent across sun elevations which result in lower inter-image variability and thus smaller JND across sun positions. This result offers the prediction that, under the global noise model, bumblebee DRA would enable them to maintain a more stable course than honeybees while flying.

Overall, our results suggest that although lowest filter-mismatch does not necessarily correspond to solar axis alignment, maximum PRC does, in agreement with the hypothesis. However, the fact that the matched filter model enables flight only towards or away from the sun suggests that complementary mechanisms must be in place to allow bees to estimate their relative position to the sun when not aligned with the solar axis. We further show that species-specific DRA anatomy shapes both the mechanism and consistency of solar axis alignment. The contribution of the fan-shaped arrangement to navigational performance likely varies across species with different visual ecologies and DRA anatomies. Fruit flies, for instance, exhibit a notably small range of ommatidial azimuth angles in their DRA^36^. This raises the question of whether the matched filter is viable for species with more restricted microvillar arrangements, a question that future computational simulations and behavioural experiments could address.

### Vector-sum model

A different, potentially complementary, mechanism to the matched filter that requires no sequential scanning of the skylight polarisation pattern and thus no active vision, is the vector-sum model^30^ (Gkanias et al., in preparation). The vector-sum model generates instantaneous sun position estimates for any body orientation by integrating PRC vectors across the DRA (Figure 3). Several behavioural experiments have shown that bees can indeed identify AoLP instantaneously^83^ while ignoring DoLP fluctuations^41^. This, however, does not exclude the possibility of parallel, sequential scanning using the matched filter.

#### Accuracy

Our results show that both the honeybee and bumblebee DRAs can estimate the sun azimuth accurately for most sun positions, with honeybees performing better on average than bumblebees (honeybees: median error=10.8°; bumblebees: median error=28.2°) (Figure 6a). This discrepancy likely arises from the viewing direction of the bumblebee DRA, as confirmed by our supplementary analyses on the forward-facing dome-shaped eye (Supplementary Figure 8). The forward-tilt causes high error estimates (∼180° error) when the sun is located behind the animal, at low and intermediate sun elevations (<45°), a known effect of a tilted DRA^30^. High error regions are rare in honeybees and limited to low sun elevations (∼0°), when the AoLP pattern is highly symmetric and when the high-DoLP band of the sky aligns more with the eye closest to the sun. This causes the eye pointing towards the anti-sun to dominate the total vector-sum (Figure 6a_*i*_). Under the vector-sum model, honeybees are predicted, thus, to orient, on average, more accurately than bumblebees.

We also summarised vector-sum performance across sun elevations by sampling each image 72 times (5° difference between each sample; total 360°) and fitting a mixture von Mises distribution to the complete set of sun position estimates. We found that the von Mises mode closest to the sun azimuth (mode_sun_) accurately captures the sun azimuth for both species (usually <10° angular deviation) (Figure 6b). The two von Mises modes are maximally separated (180° separation) when the sun is near the horizon for honeybees and for sun elevations 0°-23° for bumblebees (Figure 6c). These bimodal distributions reflect the ambiguity of sun position estimation at those elevations. It is noteworthy, that average vector-sum error at those elevations is high for both species only because of this bimodality of the total estimates, which is reflected in the low angular deviation of the mode_sun_ from the sun azimuth (Figure 6b).

Although the von Mises distributions provide a helpful summary of the vector-sum performance across sun elevations, it should be noted that any conclusions are limited by the ability of the flying animal to accurately estimate its own body rotation. More specifically, we assume that the animal can perfectly compensate for each new body orientation, i.e., each sample of the same image, to keep the sun in the same frame of reference. Overall, although bimodal von Mises distributions may reflect challenges in resolving solar and anti-solar meridians for low sun elevations, the closeness of the mode_sun_ to the true sun azimuth indicates that, despite bimodality, the correct sun azimuth is still reliably represented, potentially allowing animals to resolve this ambiguity using additional cues.

#### Consistency

The vector-sum model can be quantitatively assessed in terms of aiding bees to maintain a stable course while flying. Our analysis on vector-sum error difference (δerror) between adjacent sun positions (5° apart) show that errors are largely consistent between neighbouring sun positions (Figure 7b). This shows that small changes in sun azimuth do not, for most sun positions, elicit large changes in sun position estimation error (honeybees: median=1.6°; bumblebees: median=3.6°).

The standard deviation (SD) of the estimates across sun elevations can summarize their consistency while indicating the potential tortuosity and energetic costs of trajectories for different sun elevations (Figure 7c). For both honeybees and bumblebees, SD values are larger when the sun is at low or high sun elevations (∼180° and ∼90° SD, respectively). For the low sun elevations, the bimodality of the estimates may also lead to large SD values. As expected, overall SD values (computed over all vector-sum estimates) are generally higher than the SD of the individual von Mises modes, highlighting the potential navigational benefit of following a single mode, particularly at low sun elevations, where differences in SD are more pronounced. Thus, assuming again that animals have precise mechanical control of their rotation and the corresponding optical feedback, bees can maintain a highly stable course under the vector-sum model, mostly for intermediate sun elevations. Predictions regarding the stability of course control can be behaviourally tested by tracking flying or walking bees under different sun elevations.

## Conclusions

We provide a holistic pipeline that combines real sky images as well as empirically measured properties of honeybees’ and bumblebees’ visual systems to create a biologically accurate simulation of their polarisation vision and navigation. We assessed two different, well-described models of sun position estimation: the matched filter and the vector-sum models. The two models perform comparably both in terms of accuracy and consistency. However, the matched filter is limited to mainly identifying the anti-solar meridian only when the animal is aligned with it, while the vector-sum model performs well in estimating the sun’s azimuth from any body orientation under most sun elevations. Differences in navigational performance between the two bee species’ DRAs are mainly driven by the DRA’s viewing direction and receptive field properties.

Importantly, simulated DRAs enable direct comparative, anatomical analysis. This approach will also enable future systematic investigation of various structural and functional parameters of the visual system, unveiling evolutionary trends that may shape animals’ visual ecologies. Expanding the current framework to investigate more species’ navigational capacities and additional neural computations at later stages of visual processing will be crucial for a better understanding of the link between eye anatomy and navigation.

## Ethics

This work did not require ethical approval from a human subject or animal welfare committee.

## Data accessibility

All Data files needed to evaluate the conclusions in the paper have been uploaded to Figshare under: https://doi.org/10.6084/m9.figshare.32473422

Custom scripts used for analyses, statistical tests and visualizations have been uploaded to Figshare under: https://doi.org/10.6084/m9.figshare.32473422

## Declaration of AI use

We have not used AI-assisted technologies in creating this article.

## Authors’ contributions

G.E.K.: conceptualization, data curation, formal analysis, investigation, methodology, visualization, writing—original draft, writing—review and editing; E.G.: investigation, methodology, writing—review and editing; V.W.J.: data curation, methodology, writing—review and editing; A.A.: investigation, data curation, methodology G.C.G.: conceptualization, investigation, methodology, writing—review and editing; E.B.: investigation, methodology, writing—review and editing. B.W.: investigation, methodology, writing—review and editing. J.J.F.: conceptualization, investigation, methodology, writing—review and editing.

## Acknowledgements.

We would like to thank Daniel Calovi for his feedback on code efficiency. We would also like to thank Martín Berón de Astrada and Frida Hildebrandt for their conceptual feedback on ideas presented in this work.

